# The functional repertoire encoded within the native microbiome of the model nematode *Caenorhabditis elegans*

**DOI:** 10.1101/554345

**Authors:** Johannes Zimmermann, Nancy Obeng, Wentao Yang, Barbara Pees, Carola Petersen, Silvio Waschina, Kohar Annie Kissoyan, Jack Aidley, Marc P. Hoeppner, Boyke Bunk, Cathrin Spröer, Matthias Leippe, Katja Dierking, Christoph Kaleta, Hinrich Schulenburg

## Abstract

The microbiome is generally assumed to have a substantial influence on the biology of multicellular organisms. The exact functional contributions of the microbes are often unclear and cannot be inferred easily from 16S rRNA genotyping, which is commonly used for taxonomic characterization of the bacterial associates. In order to bridge this knowledge gap, we here analyzed the metabolic competences of the native microbiome of the model nematode *Caenorhabditis elegans*. We integrated whole genome sequences of 77 bacterial microbiome members with metabolic modelling and experimental characterization of bacterial physiology. We found that, as a community, the microbiome can synthesize all essential nutrients for *C. elegans*. Both metabolic models and experimental analyses further revealed that nutrient context can influence how bacteria interact within the microbiome. We identified key bacterial traits that are likely to influence the microbe’s ability to colonize *C. elegans* (e.g., pyruvate fermentation to acetoin) and the resulting effects on nematode fitness (e.g., hydroxyproline degradation). Considering that the microbiome is usually neglected in the comprehensive research on this nematode, the resource presented here will help our understanding of *C. elegans* biology in a more natural context. Our integrative approach moreover provides a novel, general framework to dissect microbiome-mediated functions.

## Introduction

Multicellular organisms are continuously associated with microbial communities. The ongoing interactions are likely to have influenced evolution of the involved microbes and hosts, affecting bacterial growth characteristics or host development, metabolism, immunity, and even behavior (1). Host organisms and their associated microorganisms (i.e., the microbiome) are thus widely assumed to form a functional unit, the metaorganism, where microbial traits expand host biology (2). To date, most microbiome studies focus on describing the taxonomic composition of associated communities, using 16S rRNA amplicon sequencing (3). These studies revealed that specific taxa reliably associate with certain hosts, for example Bacteroidetes and Firmicutes with humans, *Snodgrassella* and *Gilliamella* with honeybees, or *Lactobacillus* and *Acetobacter* with *Drosophila* (4–6). 16S profiling, however, is insufficient to identify bacterial functions of importance for the interaction (7). More detailed information can be obtained from bacterial genome sequences. For example, genomic analysis of the dominant members of the bee microbiome revealed complementary functions in carbohydrate metabolism, suggesting syntrophic interactions among coexisting bacteria (8). Further, the systems biology approach of constraint-based modeling permits inference of genome-scale metabolic models as a basis for predicting microbial phenotypes (9), as previously demonstrated for the interaction between whiteflies and their endosymbionts (10,11) and also hosts with more complex microbiomes (12,13).

The nematode *Caenorhabditis elegans* is one of the main model organisms in biomedical research. Yet, almost all research with this nematode has been performed in the absence of its native microbiome. In fact, its microbiome was only characterized recently, consisting mostly of Gammaproteobacteria (*Enterobacteriaceae*, *Pseudomonaceae*, and *Xanthomonodaceae*) and Bacteroidetes (*Sphingobacteriaceae*, *Weeksellaceae*, *Flavobacteriaceae*) (14–17). The little currently available data on microbiome functions highlights an influence on *C. elegans* fitness, stress resistance, and protection against pathogens (15). Previous studies also combined *C. elegans* with various soil bacteria, revealing that these can provide specific nutrients (18–22) or affect the response to drugs against cancer and diabetes (23–26). To date, the functions of the native microbiome have not yet been systematically explored.

The aim of this study was to establish the natural *C. elegans* microbiome as a model for studying microbiome functions. We extended previous 16S rRNA data (15) by sequencing whole genomes for 77 bacteria, which are associated with *C. elegans* in nature, and also *Escherichia coli* OP50, the nematode’s standard laboratory food. We reconstructed metabolic networks from the genome data to explore the metabolic competences and resulting interaction potential of the microbiome. We additionally characterized bacterial physiology and assessed which bacterial traits shape colonization ability and influence *C. elegans* fitness.

## Material and Methods

### Material

Microbiome strains were previously isolated from natural *C. elegans* isolates or corresponding substrates in Northern Germany (15; Supplementary Table S1). A representative set of 77 strains was chosen for genome sequencing. For physiological analysis, bacteria were cultured in tryptic soy broth (TSB) at 28 °C. For experiments with *C. elegans* N2, bacterial TSB cultures (500 µl at OD_600_ = 10) were spread onto peptone-free medium (PFM) agar plates. Maintenance and bleaching, to obtain gnotobiotic, age-synchronized worms, followed standard methods (27).

### Genome sequencing

Total DNA was isolated from bacterial cultures using a cetyl-trimethyl-ammonium-bromid (CTAB) approach (28). Sequencing was based on Illumina HiSeq and in a subset of nine strains additionally the PacBio platform (Supplementary Table S1). For PacBio long read genome sequencing, SMRTbell™ template library was prepared according to the manufacturer’s instructions (Pacific Biosciences, US; Protocol for Greater Than 10 kb Template Preparation). SMRT sequencing was carried out on the PacBio *RSII* (Pacific Biosciences, US) on one to three SMRT Cells, applying a movie length of 240-minutes. SMRT Cell data was assembled using the RS_HGAP_Assembly.3 protocol (SMRT Portal version 2.3.0). Chromosomes and chromids were circularized, unusual redundancies at the ends of the contigs and artificial contigs were removed after a comparison against all other replicons. Error correction was performed by Illumina reads mapping onto finished genomes using BWA (29) with subsequent variant and consensus calling using VarScan (30). QV60 consensus concordances were confirmed for all genomes. Annotations were obtained with the NCBI Prokaryotic Genome Annotation Pipeline (PGAP). For samples with only Illumina data, low quality reads and/or adaptors were trimmed with Trimmomatic v0.36 (31). *De novo* genomes were assembled using SPAdes v3.8.0 (32). Genomes (contigs greater than 1000 bp) were annotated with PGAP and Prokka v1.11 (33). Genomes were compared with BRIG (34). All sequences are available from NCBI Genbank, Bioproject PRJNA400855.

### Reconstruction of metabolic networks

Metabolic networks were reconstructed as a basis for all subsequent computational metabolic analyses and followed a two-step pipeline (Fig. 1a). First, the sequenced genomes were used to create draft metabolic models, using ModelSEED version 2.0 (35) and associated SEED reaction database. Second, we corrected errors and extended drafts by (i) finding futile cycles, (ii) allowing growth with the isolation medium (TSB), (iii) improving biosynthesis of biomass components, (iv) extending capacities to use different carbon sources, and (v) checking for additional fermentation products. This curation was based on combining topological- and sequenced-based gap filling using gapseq (version 0.9 “darwinian turtle”; https://github.com/jotech/gapseq), pathway definitions of the MetaCyc database release 22 (36), and sequence data from UniProt (37). The presence of enzymatic reactions was inferred by BLAST with bitscore of at least 50 (>=150 for a more conservative estimation), and a 75% minimum query coverage. Moreover, reactions were assumed to be present if overall pathway completeness was higher than 75% or if it was higher than 66% and key enzymes of the pathway were present (36). We also searched for genes possibly relevant in host-microbe interactions using the virulence factor database (38). The resulting curated models (Supplementary data S1) were used for further metabolic network analysis. Computations were done with GNU parallel (39).

**Fig. 1.**
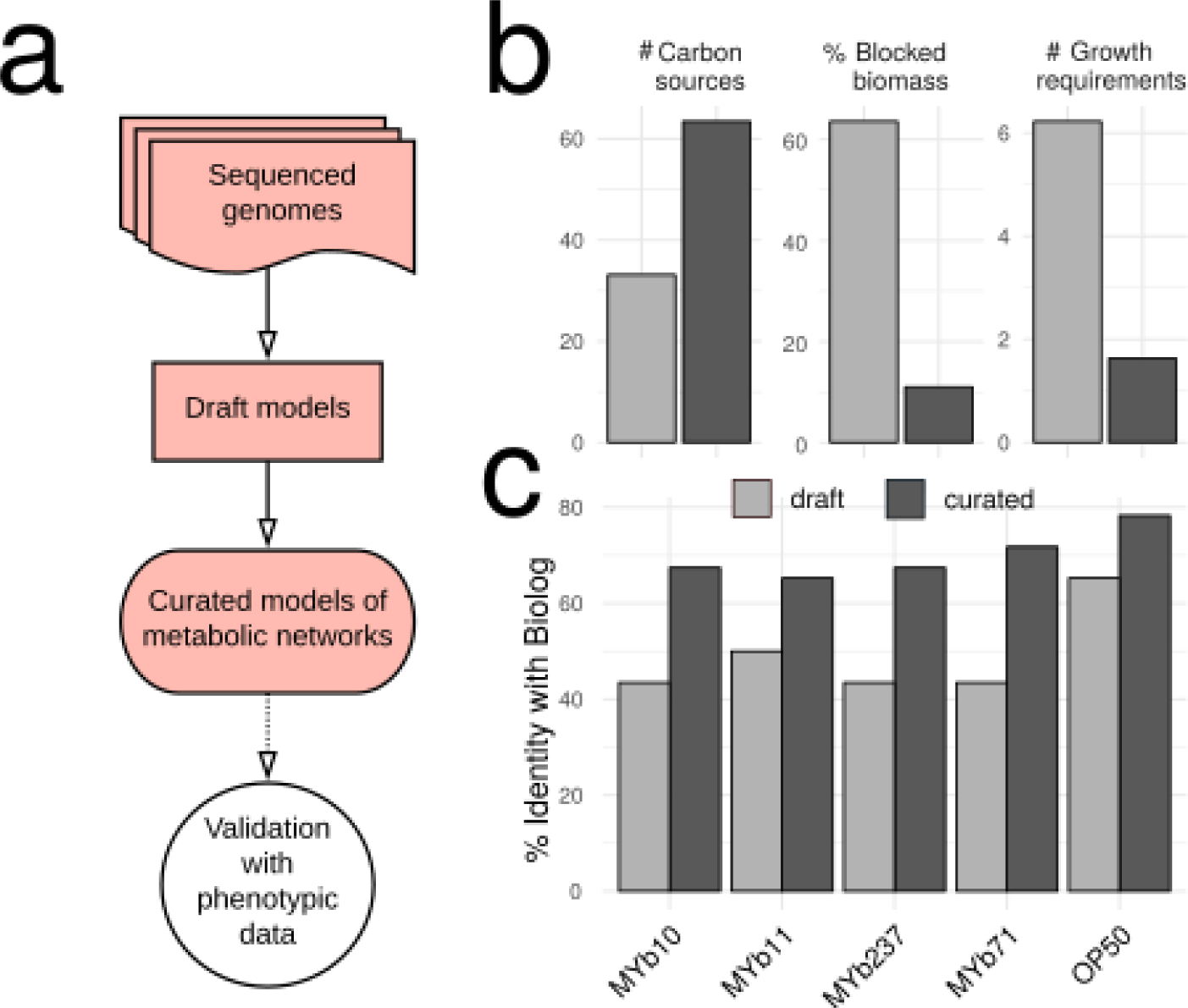
Genomes of bacterial isolates, reconstruction and validation of metabolic networks. (a) Pipeline for metabolic network reconstruction. Sequenced genomes were used to create draft metabolic models. Draft models were curated using topological- and sequenced-based gap filling. The resulting models were validated with physiological data (BIOLOG GN2; see Fig. 3); these models represent the metabolic networks of microbiome isolates and were used for functional inference. (b) Model improvements by curation, leading to an increase in accurate prediction of uptake of carbon sources, and decreases in the prediction of non-producible biomass components and the number of components needed for growth. (c) Model curation improved agreement with experimental data, as for example the BIOLOG results.

### Phylogenetic correlation and clustering of metabolic pathways

We assessed whether similarity of metabolic reactions correlated with phylogenetic relationship, using pairwise comparisons of bacteria. For each pair, the overlap of present and absent pathways (predicted by gapseq) was calculated. The corresponding 16S rRNA similarity was scored as percent identity of the global alignment using biostrings (40). 16S data was obtained from the SILVA database (41) based on the best hit of the extracted genomic 16S rRNA using RNAmmer (42). To determine overall metabolic distances between isolates, metabolic networks were treated as vectors, clustered horizontally, and metabolic distances computed as Euclidean distances between vectors. Cluster similarity was estimated by average linkage and assessed via multi-scale bootstrapping (10,000 replications) using pvclust (43).

### BIOLOG experiments

We used BIOLOG GN2 plates to assess the metabolic competence of selected bacterial strains, including MYb10, MYb11, MYb71, MYb237, and OP50. Bacterial cultures were washed three times using phosphate buffered saline (PBS) and density adjusted to OD_600_ = 1. 150 µl bacterial suspension per each well of BIOLOG plate were incubated at 28 °C for 46 h. Tetrazolium dye absorption (OD_595_) was measured every 30 min (three replicates per strain). We defined the magnitude of substrate reduction as the fold-change in tetrazolium absorbance:

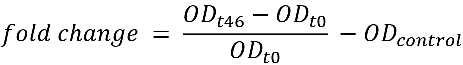

Fold-changes in water were subtracted as background. Hierarchical clustering of strains was based on average fold-change profiles (Ward’s clustering; Euclidean distance) and bootstrapping (n = 100). To analyze metabolic specialization, k-means clustering of substrates (k = 7, n = 10^3^; (44)) was performed (Supplementary Fig. S1). Statistical analyses were performed in R version 3.3.1 (45) and ggplot2 (46).

### Bacterial growth experiments

To validate BIOLOG results, we assessed growth of MYb11, MYb71, and a co-culture of both in defined media with either alpha-D-glucose or D-(+)-sucrose as carbon sources. Our defined medium is related to S medium (27), and contains 0.3% NaCl, 1 mM MgSO_4_, 1 mM CaCl_2_, 25 mM KPO_4_, 0.1% NH_4_NO_3_, 0.05 mM EDTA, 0.025 mM FeSO_4_, 0.01 mM MnCl_2_, 0.01 mM ZnSO_4_, 0.01 mM CuSO_4_, and 1% carbon source. Defined medium without carbon source served as negative and TSB as positive control. Overnight cultures were washed and adjusted to 3.94 × 10^7^ CFUs for growth experiments. Microtiter plates were incubated as BIOLOG plates above. OD_600_ was measured every 30 min, and cultures plated after 48 h. Selective plating of MYb71 using kanamycin (10 µg/ml) allowed to quantify MYb11/MYb71 proportions in co-culture. Three independent runs with technical replicates were performed and assessed with Mann-Whitney U-tests and P-value adjustment by false discovery rate (fdr).

### Simulation of bacterial in silico growth

We used the curated models to simulate the growth of MYb11 and MYb71 with sucrose as carbon source. We searched for sucrose invertases using gapseq (https://github.com/jotech/gapseq) and secreted peptides with SignalP 4.1 (47). *In silico* growth was simulated with BacArena (48). The MYb71 extracellular sucrose invertase was modeled as independent species with a single sucrose invertase reaction and exchange reactions for sucrose, glucose, and fructose. Carbon source utilization and metabolic by-products were predicted using flux balance analysis and flux variability analysis in R with sybil (49). A carbon source was assumed to be utilizable if the minimal solution of the corresponding exchange was negative (i.e., uptake) and a byproduct producible if the maximal solution of exchange positive (i.e., production).

### Simulation of ecological interactions

We assessed possible interactions among bacteria based on joined models, assuming a common compartment for metabolite exchange between microbes. Activity of individual reactions (i.e., fluxes) was linearly coupled to biomass production to prevent unrealistic exchange fluxes, such as those that solely benefit the partner but not the producer (50). The objective function was set to maximize the sum of fluxes through both biomass reactions. Two growth media were used for simulations, including TSB and a glucose minimal medium with thiamine and traces (0.001 mM) of sucrose and methionine to allow initial bacterial growth (Supplementary Table S3). Joined growth rates (j1, j2) were compared to single growth rates (s1, s2). Mutualism was defined as j1 > s1 and j2 > s2, competition as j1 < s1 and j2 < s2, parasitism as j1 < s1 and j2 > s2 (or *vice versa*), and commensalism as j1 = s1 and j2 > s2 (or the reverse).

### Experimental analysis of bacterial colonization and bacterial effects on C. elegans population growth

We examined bacterial colonization by quantifying CFUs extracted from young adult worms exposed to bacteria for 24 h. In detail, L4 larvae were placed on bacterial isolates (500 µl, OD_600_ = 10) for 24 h, washed in a series of buffers (2x M9 buffer with 25 mM tetramisole, 2x M9 with 25 mM tetramisole and 100 µg/ml gentamicin, 1x PBS with 0.025% Triton-X) to remove bacteria from the surface of nematodes, and homogenized in the GenoGrinder 2000 using 1 mm zirconia beads (1200 strokes/min, 3 min). Worm homogenate and supernatant control were plated onto TSA for quantification.

We further measured worm population growth as a proxy for worm fitness. We counted worms in the population initiated with three L4 after five days at 20 °C on bacterial lawns.

### Regression models

We analyzed the association between phenotypic measurements (i.e., bacterial colonization and worm fitness) and metabolic as well as virulence characteristics using Spearman rank correlation and random forest regression analysis. Significance for the correlation analysis was assessed with permutation tests using 100 randomly generated features and FDR-adjusted P-values. For random forest regression, the R package VSURF was used to select features based on permutation-based score of importance (51) and otherwise default settings (ntree = 2000, ntry = p/3).

### Adaptive strategies

According to the universal adaptive strategy theory (UAST) (52,53), heterotrophic bacteria follow one of three strategies: i) rapid growth and thus good competitor, ii) high resistance and thus stress-tolerator, or iii) fast niche occupation and thus ruderal. We categorized bacterial isolates using published UAST criteria (53), based on three scores, inferred from the genomes and metabolic models. In detail, the components of a competitive strategy were a large genome size, antibiotics production (presence of pathways belonging to ‘Antibiotic-Biosynthesis’ category in MetaCyc), high catabolic diversity (Metacyc: ‘Energy-Metabolism’), and siderophore biosynthesis (Metacyc: ‘Siderophores-Biosynthesis’). The criteria for stress-tolerators were auxotrophies, slow growth rates in TSB, few rRNA copies, and exopolysaccharides production (MetaCyc pathways: PWY-6773, PWY-6655, PWY-6658, PWY-1001, PWY-6068, PWY-6082, PWY-6073). The hallmarks of a ruderal strategy were fast growth in TSB, multiple rRNA copies, and low catabolic diversity (Metacyc: ‘Energy-Metabolism’). The characteristics of each isolate were related to those of the other microbiome members, yielding a relative score, thereby assuming that different strategies are present in the microbial community as a whole. For each isolate, we assessed whether the inferred value belonged to the lower or upper quantile of this criterium (in case of growth rates we used the mean instead). The total adaptive score per strategy was scaled by the number of features considered for a particular strategy. An isolate was assumed to follow the strategy, for which it produced the highest score. If two strategies had the same score, then isolates were considered to follow a mixed strategy.

## Results

### Genomes of bacterial isolates, reconstruction and validation of metabolic networks

We obtained whole genome sequences for 77 bacterial isolates of the *C. elegans* microbiome (Table 1). Of these, nine were sequenced with PacBio technology, allowing their full assembly, yielding either a single circular chromosome (four strains) or three circular chromosomes/chromids in case of the five isolates of the genus *Ochrobactrum*, which is known to have more than one chromosome (54)(Supplementary Table S1, underlined). The remaining isolates were sequenced with Illumina only, resulting in assemblies with 11 up to 243 contigs. For four genera (*Ochrobactrum*, *Pseudomonas*, *Arthrobacter*, *Microbacterium*), we included more than five strains and identified substantial intra-generic genome variation (Supplementary Fig. S2).

**Table 1.** Overview of bacterial isolates from the natural microbiome of *C. elegans* included in this study

| Phylum | Order | Genus | Isolate |
| --- | --- | --- | --- |
| Proteobacteria | Xanthomonadales | *Stenotrophomonas* | MYb238, **MYb57** |
| Proteobacteria | Pseudomonadales | *Pseudomonas* | MYb1, MYb114, MYb115, MYb117, MYb12, MYb13, MYb16, MYb17, MYb184, MYb185, MYb2, MYb22, MYb3, MYb60, MYb75, **MYb11**, MYb187, **MYb193** |
| Proteobacteria | Pseudomonadales | *Acinetobacter* | MYb10 |
| Proteobacteria | Enterobacterales | *Erwinia* | MYb121 |
| Proteobacteria | Enterobacterales | *Escherichia* | MYb137, MYb5, OP50 |
| Terrabacteria group | Actinobacteria | *Micrococcaceae* | MYb211, MYb213, MYb214, MYb216, MYb221, MYb222, MYb224, MYb227, MYb229, MYb23, MYb51 |
| Terrabacteria group | Actinobacteria | *Microbacteriaceae* | MYb24, MYb32, MYb40, MYb43, MYb45, MYb50, MYb54, MYb62, MYb64, MYb66, MYb72 |
| FCB group | Bacteroidetes | *Flavobacteriales* | MYb25, MYb44, MYb7 |
| Proteobacteria | Caulobacterales | *Brevundimonas* | MYb31, MYb33, MYb46, MYb52 |
| Terrabacteria group | Bacilli | *Paenibacillaceae* | MYb63 |
| Proteobacteria | Rhizobiales | *Ochrobactrum* | **MYb6**, MYb14, **MYb15**, MYb18, MYb19, MYb29, **MYb49**, **MYb58**, MYb68, **MYb71**, MYb237 |
| Proteobacteria | Burkholderiales | *Achromobacter* | MYb9, **MYb73** |
| Terrabacteria group | Bacilli | *Bacillaceae* | MYb48, MYb56, MYb67, MYb78, MYb209, MYb212, MYb220 |
| Bacteroidetes | Sphingobacteriales | *Sphingobacterium* | MYb181 |
| Actinobacteria | Actinomycetales | *Rhodococcus* | MYb53 |

Strains with PacBio sequencing data are given in bold.

To study the functional repertoire of the microbiome, we reconstructed genome-scale metabolic models (Fig. 1a, Supplementary data S1). The initial metabolic models were curated by screening for transporter proteins and filling of missing reactions (gap-filling). Curation increased model quality, including doubling of the number of utilized carbon sources, reduction in the absence of essential biosynthesis pathways (e.g., for nucleotides or amino acids) from 60% to below 10%, and reduction in the required additional compounds for growth on defined media from on average six to one (Fig. 1b). In order to validate our metabolic models, we experimentally quantified the ability of five selected bacterial isolates to utilize 46 carbon sources using the BIOLOG approach. The BIOLOG results produced a 49.6% overlap with the initial draft models and an increase to 70% overlap with the curated models (Fig. 1c and Supplementary Fig. S9). These curated models were subsequently used to explore bacterial metabolic competences.

### Metabolic diversity within the microbiome of C. elegans

Using the metabolic networks, we assessed a possible relationship between metabolic and phylogenetic similarities and explored the metabolic potential of the isolates. We found that the information contained in pairwise 16S rRNA phylogenetic relationships is generally indicative of the corresponding similarities in metabolic networks (Fig. 2a; Spearman rank correlation, *R_S_* = 0.6199, *P* < 0.0001). Metabolic similarities appeared to be higher than phylogenetic relationships, suggesting a considerable overlap in metabolic competences across the included isolates. Nevertheless, some variation was identified, even among isolates from the same genus. Such variation within taxonomic groups was confirmed through hierarchical clustering of the inferred metabolic networks (Fig. 2b), as for example seen for the *Pseudomonas* isolates, which contain three clearly separated clusters. Similar patterns are also observed for other genera, for example *Enterobacter*, *Ochrobactrum*, or *Microbacterium*. We conclude that variation in metabolic competences is generally related to the bacterial phylogeny albeit some variation being present within genera.

**Fig. 2.**
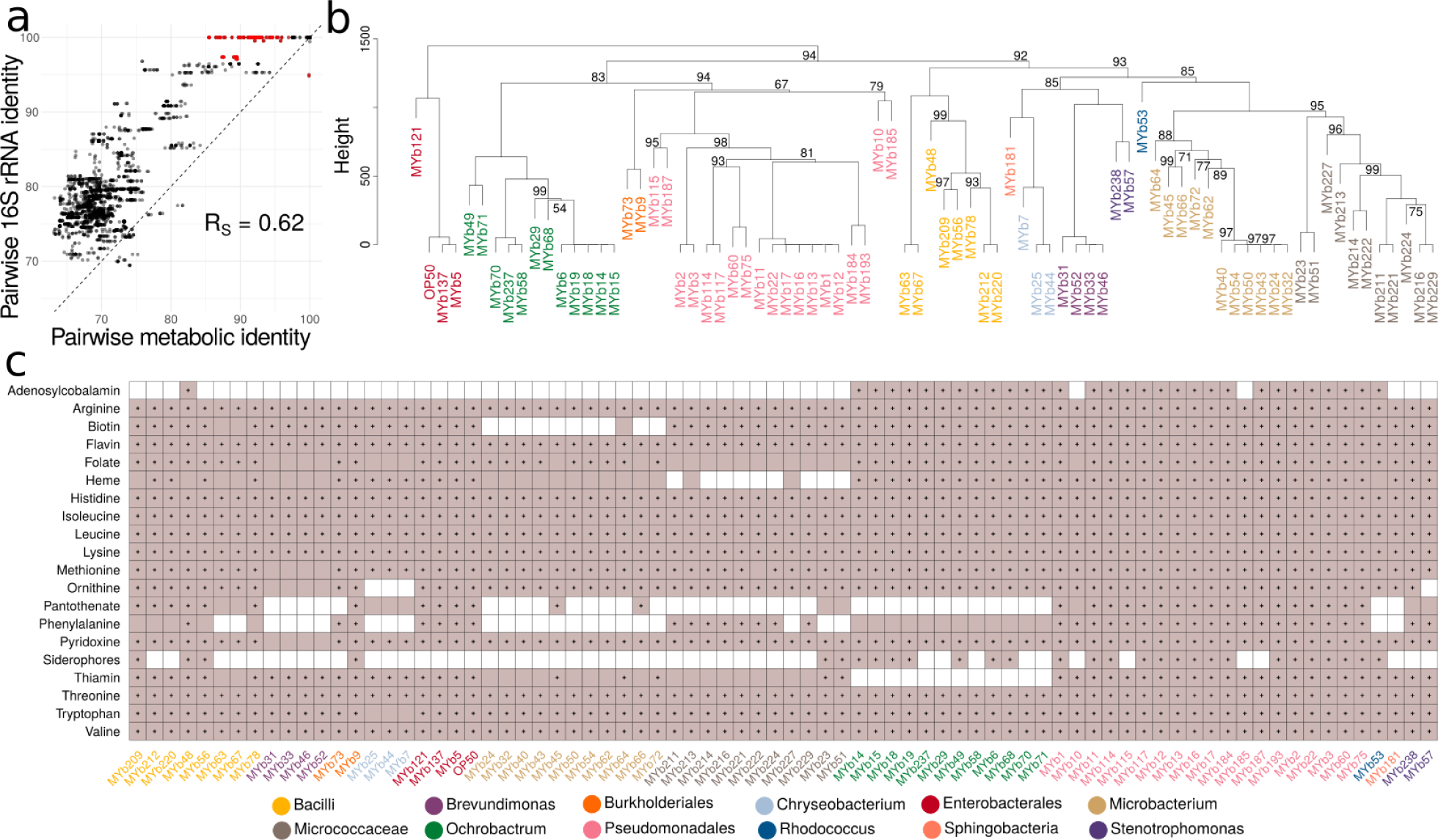
Metabolic network clustering and distribution of important pathways. (a) Correlation between pairwise similarities in 16S rRNA sequences and metabolic networks is shown. Red indicates pairs with a 16S rRNA identity above 97% and metabolic identity below 97% and vice versa. (b) Hierarchical clustering of metabolic networks based on pathway prediction. P-values were calculated via multiscale bootstrap resampling. In case of full support (i.e., *P* = 100), P-values are not shown (For a complete list of different unbiased P-values and bootstrap values see Supplementary Figure S11). (c) Prediction of bacterial capacity to produce metabolites favoring *C. elegans* growth. Filled squares in light purple indicate that the metabolic networks predict presence of the biosynthetic pathway required to produce essential amino acids and co-factors. Black dots within the filled squares indicate that pathway presence is supported by more conservative parameters (BLAST bit score >= 150). Different bacterial genera in (b) and (c) are indicated by different colors of the strain names (Table 1).

We next assessed the metabolic competences of the microbiome isolates (Supplementary Table S4). In general, the inferred metabolic competences are consistent with the aerobic and heterotrophic lifestyle of the *C. elegans* host. The glycolysis, at least the partial pentose phosphate pathway, the tricarboxylic acid cycle, and enzymes enabling oxidative phosphorylation (cytochrome oxidases) were present in all genomes. Almost all isolates possessed enzymes enabling tolerance to microaerobic conditions (e.g., cytochrome bd oxidase). Some isolates from Bacilli, *Pseudomonas*, and *Ochrobactrum* showed sequence-evidence for chemolithotrophic life style (nitrite and formate oxidation) and anaerobic respiration (nitrate, arsenate reduction). Pathways related to CO_2_ fixation (reductive TCA or anaplerosis) were found in a few *Pseudomonas*, Bacilli, or *Microbacterium* isolates. Two Bacillales strains further showed capacity to degrade polysaccharides, such as starch, cellulose, mannan, rhamnogalacturonan (e.g., *Paenibacillus* MYb63, *Bacillus* MYb67). The microbiome members are able to produce all essential substances required for *C. elegans* growth, which the nematode cannot synthesize on its own (i.e., all essential amino acids and vitamins; Fig. 2c). Most variation among isolates was observed in the biosynthetic pathways of B12, pantothenate, phenylalanine and siderophores (Fig. 2c). Simulation of *in silico* growth (Supplementary Fig. S9) suggests that simple sugars, such as glucose, ribose or arabinose, can be used by all organisms while the ability to degrade lactose, maltodextrin, or sucrose varies among strains. Short chain fatty acids were among the compounds that can be generated by all organisms (Supplementary Fig. S9), while there was variation in the ability to produce succinate, cysteine, and valine. Moreover, several microbiome members possessed potential virulence genes, especially the *Pseudomonas* and *Escherichia* isolates (Supplementary Table S5).

We subsequently focused our analysis on *Ochrobactrum* and *Pseudomonas* isolates. These two genera are enriched in the native microbiome of *C. elegans*, comprising 10– 20 % of the associated bacteria, they are also particularly well able to colonize the nematode gut (15), and some isolates can protect *C. elegans* from pathogen infection (15,55). Most *Pseudomonas* isolates can provide all required substances for nematode growth. *Ochrobactrum* isolates are able to produce vitamin B12, like *Pseudomonas* isolates, but unlike almost any of the other microbiome members (Fig. 2c). Moreover, the *Ochrobactrum* isolates vary from other microbiome members in degradation pathways, energy metabolism, vitamin biosynthesis, and presence of potential virulence factors (Supplementary Table S6). These isolates appear to lack some for *C. elegans* relevant vitamin biosynthetic pathways, such as those leading to thiamine and panthothenate. They possess a unique Brucella-like putatively immune-modulating LPS (Supplementary Table S5).

In summary, we found that *C. elegans* harbors a microbial community with diverse metabolic competences, which can supply all essential nutrients for *C. elegans* and which includes several *Ochrobactrum* and *Pseudomonas* isolates capable of producing important vitamins such as vitamin B12.

### Nutrient context influences ecological interactions within the microbiome

To study how metabolic repertoires affect bacterial growth and interactions within the microbiome, we characterized carbon source utilization of selected isolates and tested growth in different nutrient environments *in vitro* and *in silico*. Using the BIOLOG approach, we focused on prominent *C. elegans* microbiome members that colonize worms and affect host fitness, including *Ochrobactrum* sp. MYb71, *Ochrobactrum* sp. MYb237, *Acinetobacter* sp. MYb10, *Pseudomonas lurida* MYb11, and as a contrast the laboratory food strain *E. coli* OP50 (Supplementary Fig. S3; (15)). For a first insight into bacterial interactions, we additionally included a MYb11-MYb71 mixture (two strains that can co-exist in *C. elegans* (15). We found that the metabolic repertoires of the strains differ and that the four microbiome isolates can be distinguished from OP50 based on the metabolism of carboxylic and amino acids (Fig. 3a, cluster II; Supplementary Fig. S4). Within the microbiome, MYb10 was least versatile at using carboxylic acids and sugar alcohols (Fig. 3a, cluster IV), while MYb11 and the two *Ochrobactrum* strains could additionally metabolize unique sets of carboxylic acids and sugar alcohols, respectively (Fig. 3a, cluster V and III). Notably, the disaccharides sucrose and turanose were only metabolized by MYb71 (alone and in co-culture) (Fig. 3a, cluster III), although sucrose invertases were present in the genomes of both MYb71 and MYb11 (cf. pathway: sucrose degradation I, Supplementary Table S4). In co-culture, the metabolic repertoires of MYb11 and MYb71 appeared additive.

**Fig. 3.**
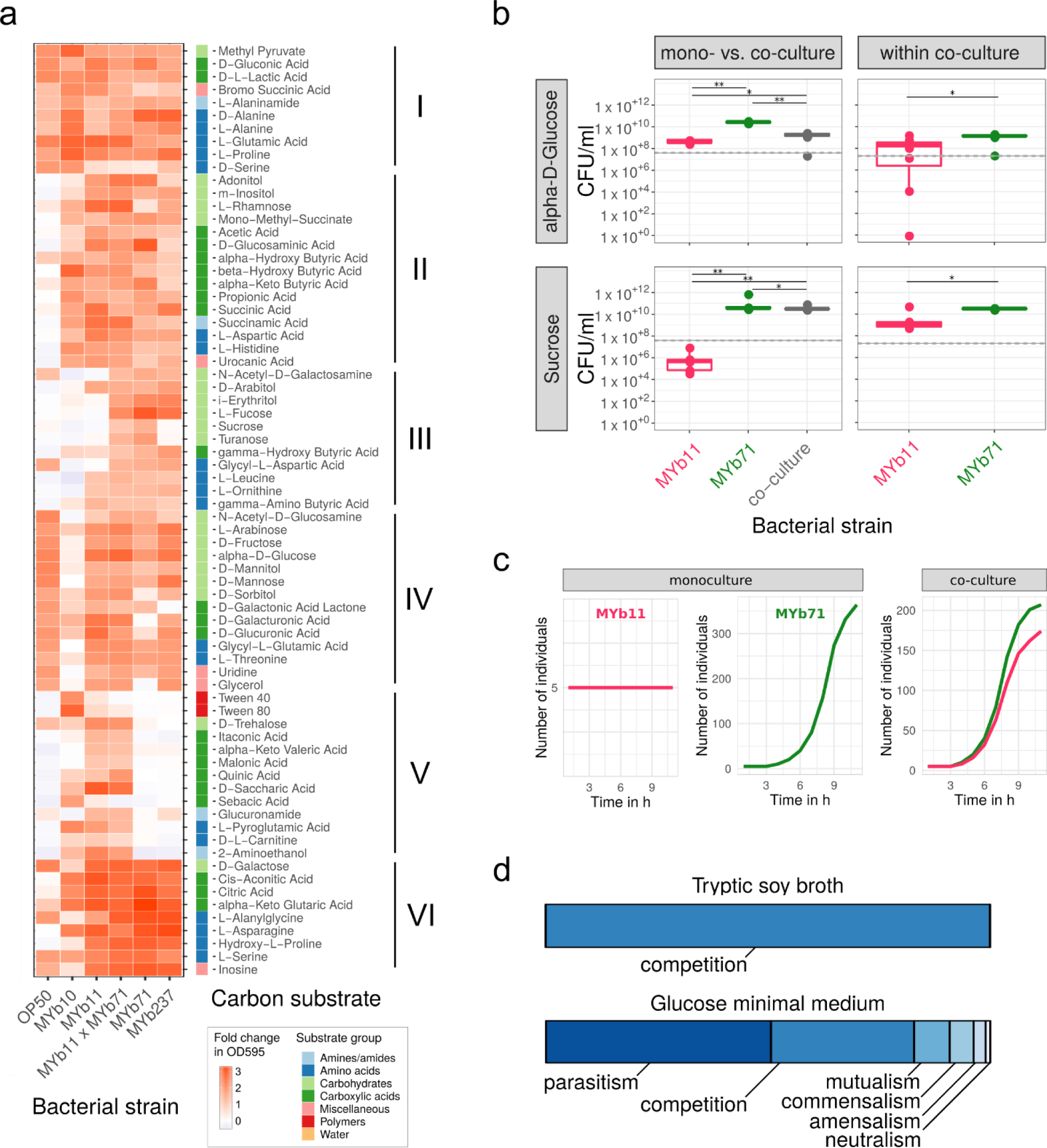
Realized carbon metabolism and growth. (a) Profiles of carbon substrate use of *Acinetobacter* sp. (MYb10), *Pseudomonas lurida* (MYb11), *Ochrobactrum* sp. (MYb71), *Ochrobactrum* sp. (MYb237), and *E. coli* OP50 in BIOLOG GN2 plates over 46 h. The fold-change in indicator dye absorption from 0 to 46 h indicates that the particular compound is metabolized. K-means clustering (k = 7) of substrates by fold-change highlights metabolic differences between strains. See Supplementary Fig. S5 for cluster VII with substrates used poorly across most strains. (b) Colony forming units per ml (CFU/ml) of MYb11 and MYb71 in mono- and co-culture at 48 h in alpha-D-glucose and sucrose-containing minimal media. The horizontal and dashed lines indicate mean and SD of CFU/ml at inoculation. Statistical differences were determined using Mann-Whitney U-tests and corrected for multiple testing using fdr, where appropriate. Significant differences are indicated by stars (** for *p* < 0.01; * for *p* < 0.05). Data from three independent experiments is shown. (c) *In silico* growth of MYb11 and MYb71 in mono- and co-culture in sucrose-thiamine medium using BacArena with an arena of 20×20 and five initial cells per species. (d) Bacterial interaction types observed during *in silico* co-cultures of all combinations of the 77 microbiome isolates and OP50.

We next assessed whether the differences in MYb11 and MYb71 metabolic competences shape bacterial interactions in growth media with only a single carbon source. We did not observe any growth in a control medium without a carbon source, and thus conclude that the tested bacteria are not chemoautotrophic (Supplementary Fig. S6). In minimal medium with alpha-D-glucose, both MYb11 and MYb71 grew, yet exhibited distinct growth dynamics (Fig. 3b; Supplementary Fig. S6). MYb71 produced more CFUs than MYb11 in co-culture (Fig. 3b), suggesting that MYb71 has a growth advantage over MYb11 and/or interferes with MYb11 in some other way. In agreement with the BIOLOG results, a medium including sucrose as the sole carbon source supported only growth of MYb71 but not MYb11 in monoculture (Fig. 3b, Supplementary Fig. S6). Surprisingly, MYb11 increased in CFUs in co-culture, while in monoculture MYb11 CFUs declined over time, indicating parasitic growth (Fig. 3b). Thus, the presence of different carbon sources can change the interaction type between two isolates.

We subsequently analysed in more detail the basis for co-growth of MYb11 and MYb71 in sucrose medium, using genome sequence information and *in silico* growth simulations. Interestingly, we found a secreted sucrose invertase in the genome of MYb71 but not MYb11 (Supplementary Fig. S10). *In silico* growth simulations demonstrated that MYb71 can grow in sucrose medium, MYb11 alone does not, while in co-culture both increased in numbers (Fig. 3c). The simulations thereby re-captured the *in vitro* findings of distinct growth patterns in sucrose medium. Genome sequence information strongly suggests that growth of both in co-culture is mediated by a secreted enzyme from MYb71.

Taking a more global perspective, we next investigated *in silico* the potential ecological interactions among the 77 microbiome isolates and *E. coli* OP50. We compared the growth characteristics of single bacteria with co-growth rates of all 3003 possible bacterial pairs in different nutrient environments. In a rich medium (TSB), the exclusive interaction type was competition, indicated by lower growth rates in co-vs. mono-culture (Fig. 3d). This changed completely when the nutrient environment was simplified to a glucose minimal medium: 50% of the interactions were parasitic (i.e., the growth rate for one isolate was higher in co-culture than in monoculture, while this pattern was opposite for the other isolate of a pair), one out of three interactions were competitive, and 8% mutualistic (i.e., growth rates for both isolates higher in co-culture than the monocultures; Fig. 3d). Under these minimal medium conditions, the most frequently exchanged metabolites across bacteria were glyceraldehyde, acetate, and ethanol (Supplementary Fig. S7). We conclude that the nutrient context modulates bacterial growth, consistently identified both *in silico* and *in vitro*, and thereby shapes bacteria-bacteria interactions within the microbiome.

### Specific metabolic competences predict bacterial colonization ability and bacterial effects on nematode fitness

To characterize traits involved in the interaction between bacteria and *C. elegans*, we identified genomic and metabolic features that are associated with the bacteria’s colonization ability and their effects on worm fitness. We focused on 18 microbiome isolates based on (i) their abundance in the *C. elegans* microbiome, (ii) enrichment in worms, and (iii) effects on worm population growth (15,56). OP50 was included as control. Our phenotypic analysis revealed substantial variation among bacterial isolates in both their ability to colonize *C. elegans* and their effects on nematode fitness (Fig. 4; Supplementary Fig. S3). Importantly, these two microbiome characteristics were significantly related with certain metabolic competences of the bacteria. Pyruvate fermentation to (S)-acetoin was significantly associated with bacterial load and the degradation of trans-3-hydroxyproline with nematode population growth (Fig. 4, Supplementary Table S8).

**Fig. 4.**
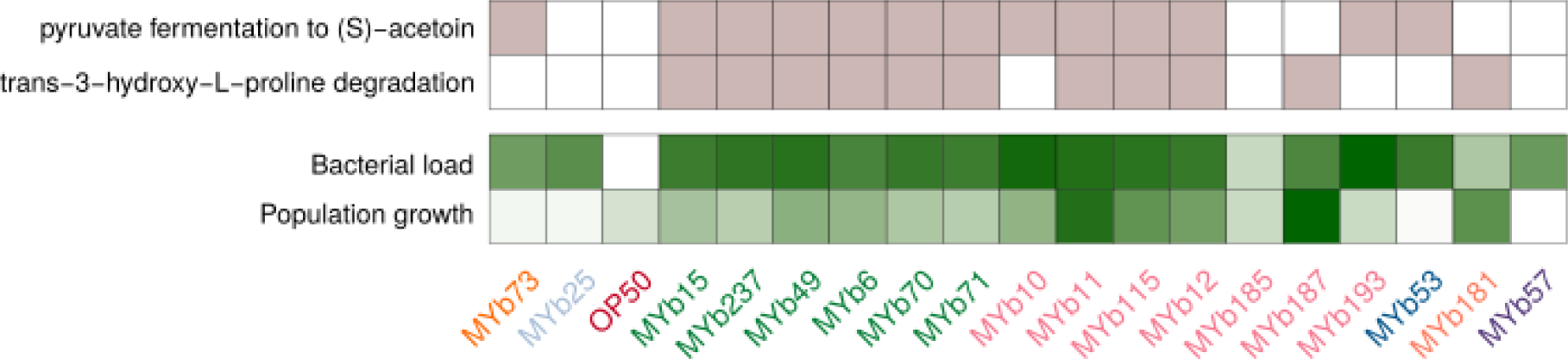
Relationship of bacterial metabolic competences with their colonization ability and their effects on nematode fitness. Presence of metabolic traits (light purple color), which were found to be associated with the bacteria’s ability to colonize *C. elegans* or affect nematode population growth as a proxy for worm fitness (green color). Regression models suggested that the pathway of pyruvate fermentation to acetoin influences bacterial load while the presence of hydroxyproline degradation is associated with *C. elegans* population growth. Colonization and population growth data was normalized; darker colors indicate increased capacities. Different bacterial genera are indicated by the different colors of the strain names (Table 1).

To further explore the potential behavior of the microbiome isolates in an ecological context, we interpreted their genomic and metabolic traits in light of the universal adaptive strategy theory (52,53). We found that 26 isolates were associated with a competitive, 9 with a stress-tolerating, and 37 with a ruderal (fast niche occupiers) strategy (Fig. 5a). The remaining 6 isolates showed a mixed strategy (same score for competition and stress-tolerance). Interestingly, bacterial isolates with different adaptive strategies also varied in their colonization ability (Fig. 5b): Bacterial isolates with competitive or stress-tolerance strategies showed higher bacterial load in *C. elegans* than those with ruderal strategy (Wilcoxon rank sum test, P = 0.01). Moreover, for the competitive and stress-tolerance isolates, we identified a positive correlation between bacterial load and the inferred score (Spearman, R_S_ = 0.37, *P* = 0.1; Supplementary Fig. S8). Taken together, the competitive and stress-tolerating strategies are most prevalent within the microbiome of *C. elegans* and relate to bacterial colonization capacity.

**Fig. 5.**
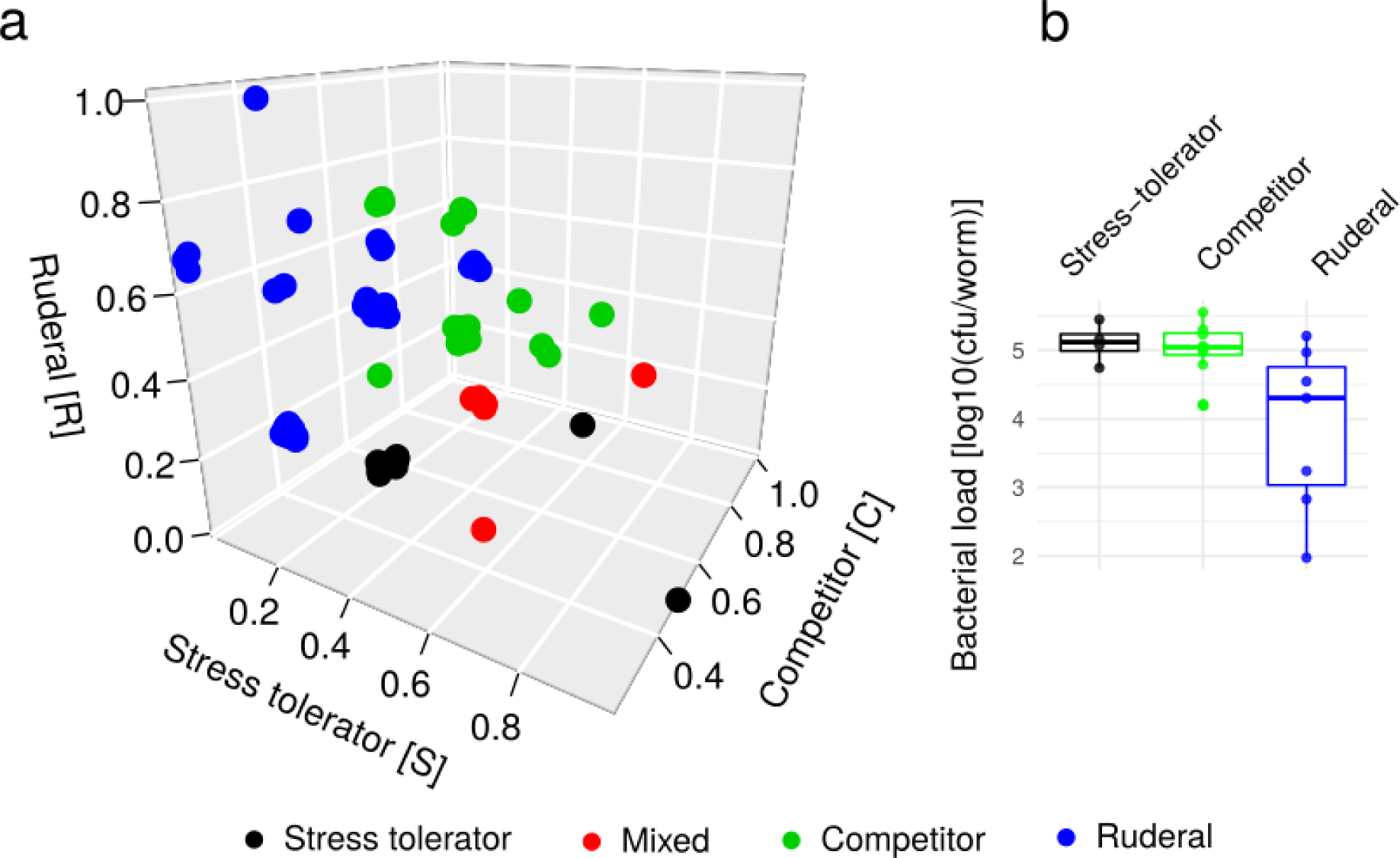
Different adaptive strategies within the microbiome and their relationship to worm colonization. We applied the universal adaptive strategy theory proposed for soil bacteria (48) to categorize the bacterial isolates. (a) Based on genomic and metabolic features, each isolate obtained a score for the competitive (C), stress tolerating (S), and ruderal (R) strategy, which is represented in the 3D-coordinate system. (b) Bacterial colonization behavior in comparison to adaptive strategies. Isolates which were categorized as mixed strategists (i.e. same score for competitive and stress-tolerance) produced the lowest bacterial load, whereas stress-tolerator and competitors had the highest values. The difference in bacterial load between ruderal and other strategies was significant (Wilcoxon rank sum test, *P* = 0.01).

## Discussion

We here present the first overview of the functional repertoire encoded within the native microbiome of the model organism *C. elegans* and provide a metabolic framework for the functional analysis of host-associated microbial communities, which combines genome sequence data, metabolic network modeling, and physiological characterizations. For our analysis, we obtained whole genome sequences and reconstructed the metabolic network of 77 microbiome members. We identified variation in the metabolic competences within the microbiome and found that the community as a whole is able to produce nutrients essential for *C. elegans* growth. For selected bacteria, we were able to validate the model predictions with the help of physiological analyses. Moreover, we used both *in vitro* and *in silico* approaches to demonstrate that the nutrient environment can lead to a shift in the interaction between bacteria, for example from competition to mutualism. We further identified specific metabolic modules that appear to shape the interaction with the nematode host, including pyruvate fermentation to (S)-acetoin and the degradation of trans-3-hydroxyproline. Finally, we considered a combination of genomic, metabolic and cellular traits to infer bacterial life history strategies according to the universal adaptive strategy theory (52,53), finding that bacterial colonization ability is associated with a competitive or stress-tolerant strategy. In the following, we will discuss in more detail (i) the diversity of metabolic competences in the microbiome and possible implications for *C. elegans* biology, (ii) how the metabolic networks shape bacteria-bacteria interactions, and (iii) in which ways bacterial traits can affect colonization and *C. elegans* fitness.

Our analysis revealed that the microbiome members are jointly able to synthesize all essential nutrients required by *C. elegans*. Individual bacterial isolates are not able to do so. The considered isolates varied in their capacity to produce vitamins essential to *C. elegans*, such as folate, thiamine, and vitamin B12, which are known to affect nematode physiology and life history (20,21,23,57–59). For example, vitamin B12 influences propionate breakdown, it can accelerate development, and reduce fertility (58,59). Within the microbiome isolates studied here, only *Pseudomonas* and *Ochrobactrum* strains had the pathways to produce vitamin B12. An enrichment of these genera in the microbiome should therefore affect both the metabolic state and fitness of *C. elegans*.

Our study demonstrated that bacterial interactions can change depending on the nutrient environment. In our simulations, competitive interactions dominated in rich medium (TSB), while parasitic and mutualistic interactions in minimal medium. Furthermore, interactions between *Pseudomonas lurida* (MYb11) and *Ochrobactrum* sp. (MYb71) shifted from parasitic to competitive in a sucrose- vs. glucose-supplemented medium. In line with this, we have detected a secreted sucrose invertase in the genome of MYb71, which otherwise lacks any known sucrose transporter. Thus, we propose that MYb71 breaks down sucrose extracellularly, and the monosaccharides glucose and fructose become exploitable by MYb11. While a similar phenomenon has been described in cultures of yeast with engineered auxotrophies (60,61), we here observed this capacity in strains from a natural community of host-associated microbes. This emphasizes the relevance of nutrient context in host-microbiome interactions. Importantly, no single growth medium might reliably predict all possible interaction types among bacteria. It is therefore essential to consider the nutrient context in order to fully understand the range of possible interactions within the microbiome (e.g. (62)).

Our analysis further identified two bacterial traits that appear to influence colonization ability and impact *C. elegans* fitness. Colonization ability was associated with pyruvate fermentation to (S)-acetoin. This fermentation pathway includes the ketone diacetyl as an intermediate, whose buttery odor attracts *C. elegans* and promotes feeding behaviour (63). In detail, diacetyl binds the transmembrane odor receptor *odr-10* and induces a shift in odortaxis (63–65). As a result, worms are more attracted to and less likely to leave bacterial lawns with this particular smell (63). Indeed, lactic acid bacteria in rotting citrus fruits were more attractive to worms when releasing diacetyl (66). Similarly, entomopathogenic nematodes of the genus *Steinernema* were more attracted to insect cadavers, when they were infected with the diacetyl-producing bacterial symbionts of the nematode (67). Thus, if worms are attracted to diacetyl-producing bacteria, they should spend more time in their presence. This alone could increase uptake of bacteria and subsequently bacterial colonization.

We also found that trans-3-hydroxyproline degradation in bacteria is associated with increased nematode fitness. In worms, hydroxyproline is present in collagen type IV, a major component of the extracellular matrix in the pharynx, intestine and cuticle (68–70). The breakdown of hydroxyproline can generate reactive oxygen species (71). These may act as signaling molecules, which could affect cellular proliferation (72) and *C. elegans* reproduction (73). Whether ROS in the gut increases brood size is unknown. Alternatively, bacteria with the degradation pathway may utilize the amino acid as a carbon source, consistent with the “microbiome on the leash” hypothesis, characterized by host-selection of bacterial traits (74) and where nematodes indirectly benefit, if damage is limited and if it allows worms to maintain a beneficial symbiont.

Our analysis of bacterial life history strategies (52,53) additionally identified competitiveness and stress tolerance to associate with *C. elegans* colonization. It appears that the ability to outcompete other microbes and/or tolerate stress environments is an important requirement for a close relationship with the nematode. By linking bacterial genomics, metabolism, and physiology with worm phenotypes, we were thus able to generate testable hypotheses on traits and adaptive strategies important for life in the worm.

In conclusion, our study provides a resource of naturally associated bacteria, their whole genome sequences, and reconstructed metabolic competences that can be exploited to study and understand *C. elegans* in an ecologically meaningful context. This resource may help to further establish *C. elegans* as a versatile model for studying the genetics of host-microbe interactions. It may also prove useful to characterize a variety of phenotypes and underlying molecular mechanisms in *C. elegans*, which have thus far only been studied in the complete absence of the worm’s microbiome.

## Acknowledgements

We thank Simone Severitt and Nicole Heyer for technical assistance regarding PacBio genome sequencing, Jolantha Swiderski for long read genome assemblies, Peter Deines for advice on BIOLOG assays, and the BiMo/LMB of Kiel University for access to their core facilities. We are grateful for funding from the Deutsche Forschungsgemeinschaft (DFG, German Research Foundation) within the Collaborative Research Center CRC 1182 on Origin and Function of Metaorganisms, projects A1 (HS, KD, ML), A4 (HS), and INF (MPH, CK). We further acknowledge funding by the DFG under Germany’s Excellence Strategy – EXC 2167-390884018 (Excellence Cluster Precision Medicine in Chronic Inflammation; CK, HS), the Competence Center for Genome Analysis Kiel (CCGA Kiel; HS), the Max-Planck Society (Fellowship to HS), and the International Max-Planck Research School for Evolutionary Biology (NO).

## Conflict of Interest

The authors declare no conflict of interest.

Supplementary information is available at ISME’s website.

## Notes

**Conflict of interest**: All authors declare no competing financial interests in relation to the work described.

