## Supplementary File for "The functional repertoire encoded within the native microbiome of the model nematode *Caenorhabditis elegans*"

#### Title

#### Running title

*C. elegans* microbiome functions

#### Authors

Johannes Zimmermann[1]*, Nancy Obeng[2]*, WentaoYang[2], Barbara Pees[3], Carola Petersen[2,3], Silvio Waschina[1], Kohar Annie Kissoyan[2], Jack Aidley[2], Marc P. Hoeppner[4], Boyke Bunk[5], Cathrin Spröer[5], Matthias Leippe[3], Katja Dierking[2], Christoph Kaleta[1#], Hinrich Schulenburg[2,6#]

#### Affiliations

1 Research Group Medical Systems Biology, Institute of Experimental Medicine, Christian-Albrechts University, Kiel, Germany

2 Research Group of Evolutionary Ecology and Genetics, Zoological Institute, Christian-Albrechts University, Kiel, Germany

3 Research Group of Comparative Immunobiology, Zoological Institute, Christian-Albrechts University, Kiel, Germany

4 Institute of Clinical Molecular Biology, Christian-Albrechts University, Kiel, Germany

5 Leibniz Institute DSMZ-German Collection of Microorganisms and Cell Cultures, Braunschweig, Germany

6 Max-Planck Institute for Evolutionary Biology, Ploen, Germany

* These authors contributed equally to this work: Shared first authorship

### These authors contributed equally to this work: Shared senior authorship

**Correspondence**

Christoph Kaleta, Research Group Medical Systems Biology, Institute of Experimental Medicine, Christian-Albrechts University, Michaelisstraße 5, 24105 Kiel, Germany; Tel: +49-431-50030340; Fax: +49-431-50030344;

Hinrich Schulenburg, Research Group of Evolutionary Ecology and Genetics, Zoological Institute, Christian-Albrechts University, Am Botanischen Garten 9, 24118 Kiel, Germany; Tel.: +49-431-8804141; Fax: +49-431-8802403;

**Conflict of interest**

All authors declare no competing financial interests in relation to the work described.

**Keywords**

Microbiome, Microbiota, *Caenorhabditis elegans*, *Ochrobactrum*, *Pseudomonas*, Metabolic networks

Financial support: German Science Foundation Collaborative Research Center CRC 1182 on Origin and Function of Metaorganisms, projects A1 (KD, ML, HS), A4 (HS), and INF (MPH, CK). Excellence Cluster Precision Medicine in Chronic Inflammation (PMI; CK, HS); the Competence Center for Genome Analysis Kiel (CCGA Kiel; HS); the Max-Planck Society (Fellowship to HS); and the International Max-Planck Research School for Evolutionary Biology (NO).

#### Supplementary materials

*Supplementary figures*

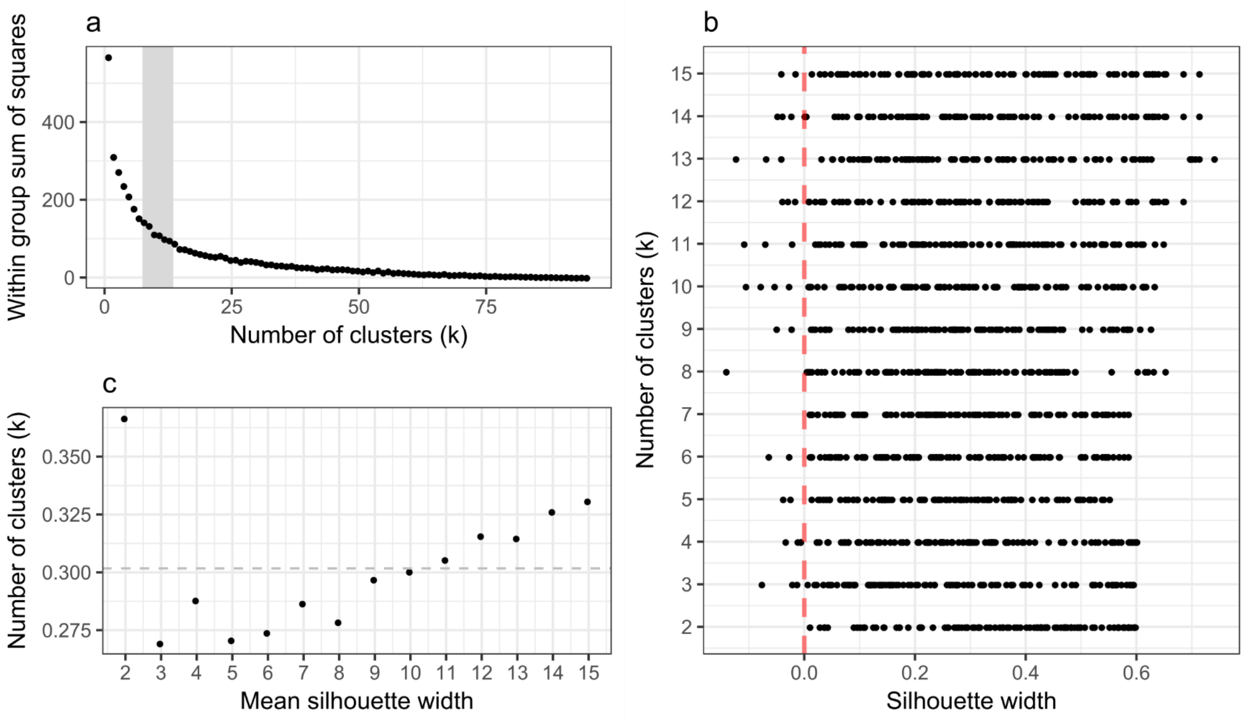


**Supplementary Fig. S1. Diagnostic plots of k-means clustering.** (a) Within group sum of squares (SS) across clustering with different k. Grey shaded region highlights number of clusters where within group SS starts decreasing less rapidly with an increase in k. (b) Silhouette width of clustering with different k. A silhouette width = 0 indicates points of several clusters overlapping, while a silhouette width = 1 implies a point being exclusive to one cluster. (c) Mean silhouette width across clustering with different k 100 times. Dashed grey line indicates overall mean silhouette width observed.

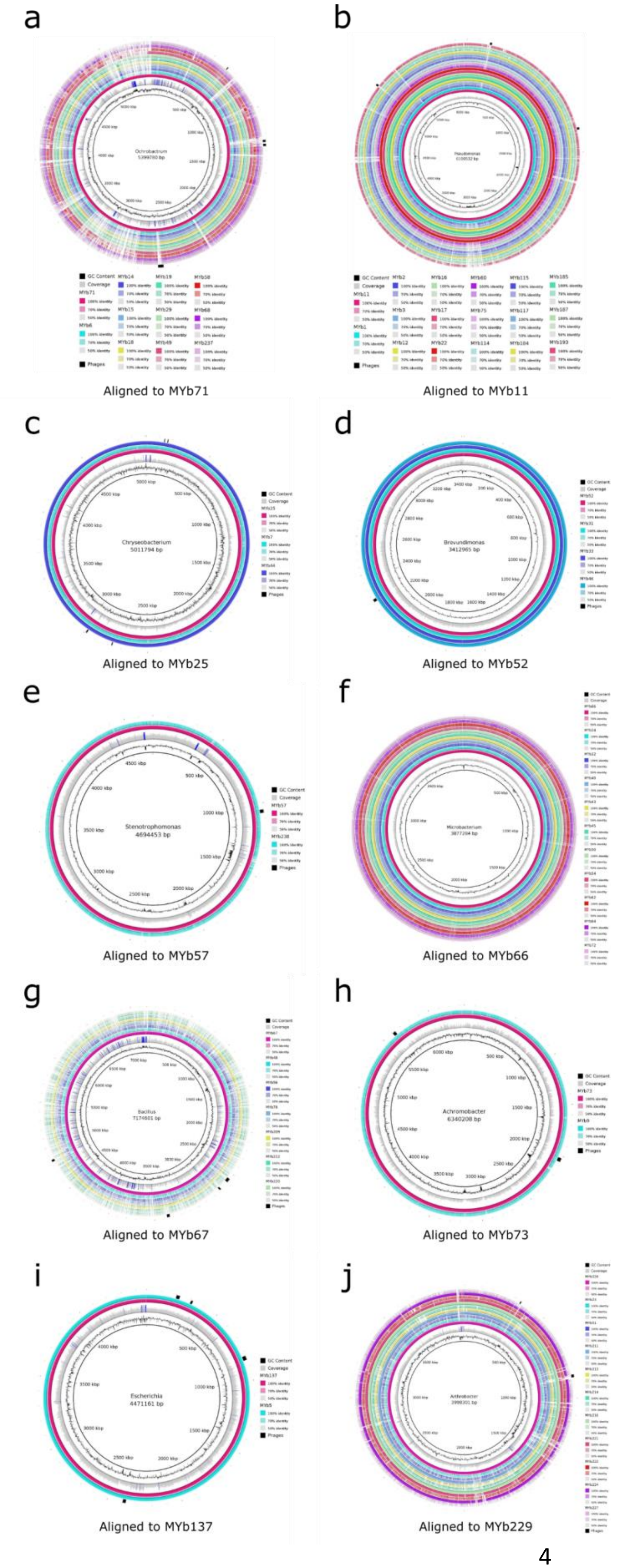
a
Aligned to MYb71
b
Aligned to MYb11
c
Aligned to MYb25
d
Aligned to MYb52
e
Aligned to MYb57
f
Aligned to MYb66
g
Aligned to MYb67
h
Aligned to MYb73
i
Aligned to MYb137
j
Aligned to MYb229

**Supplementary Fig. S2. Genome comparison of selected bacterial taxa using circular plots.** Selected taxa include: (a) *Ochrobactrum*, (b) *Pseudomonas,* (c) *Chryseobacterium*, (d) *Brevundimonas*, (e) *Stenotrophomonas*, (f) *Microbacterium*, (g) *Bacillus*, (h) *Achromobacter*, (i) *Escherichia*, and (j) *Arthrobacter*. For each taxon, the genome sequence with highest quality (i.e., genome sequence with the smallest number of contigs) was identified and then used as a reference for alignment of the remaining genomes. GC content, coverage, and predicted bacterial phage information are given for the reference genome of each taxon. The color intensity in each ring indicates BLAST match identity. The genomes within a particular taxon are usually highly similar, except in the case of *Bacillus*, which produces higher diversity across the included isolates. Predicted phage regions tend to be highly diverse across isolates within a taxon, except for *Chryseobacterium*, *Brevundimonas*, *Stenotrophomonas*, *Achromobacter*, and *Escherichia*, which may be due to the small number of species in these taxa.

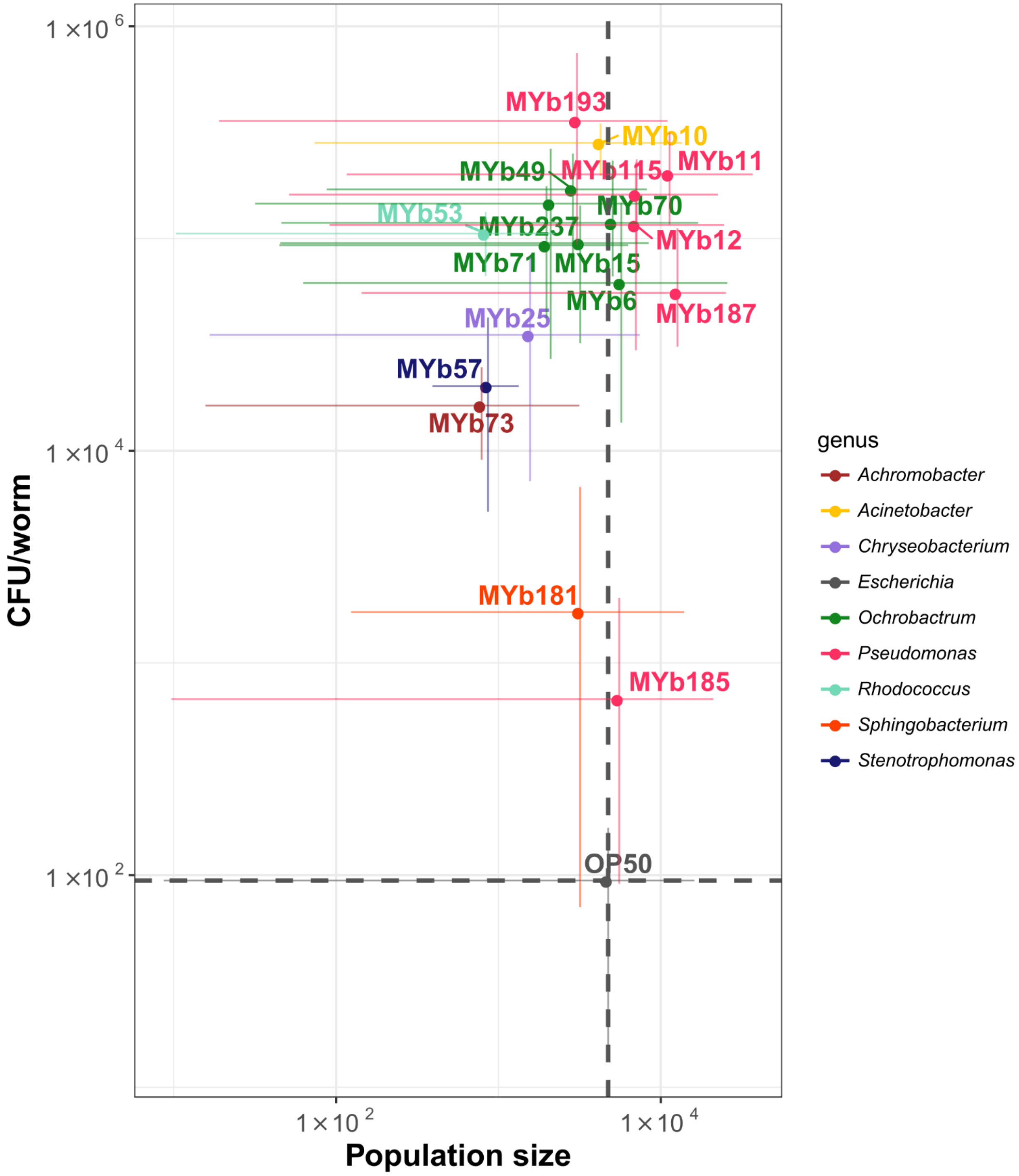


**Supplementary Fig S3. Variation in population growth and colonization levels of *C. elegans* in mono-association with natural microbiome isolates.** Three L4 larvae were exposed to bacterial lawns on PFM plates for five days, and F2 population sizes quantified (n = 3-6). To count colony forming units per worm, L4 larvae were transferred from NGM plates with OP50, exposed to microbiota lawns for 24 h, washed and the associated bacteria extracted (n = 5). Dashed lines show the mean population size and bacterial load of the canonical food bacteria *E. coli* OP50.

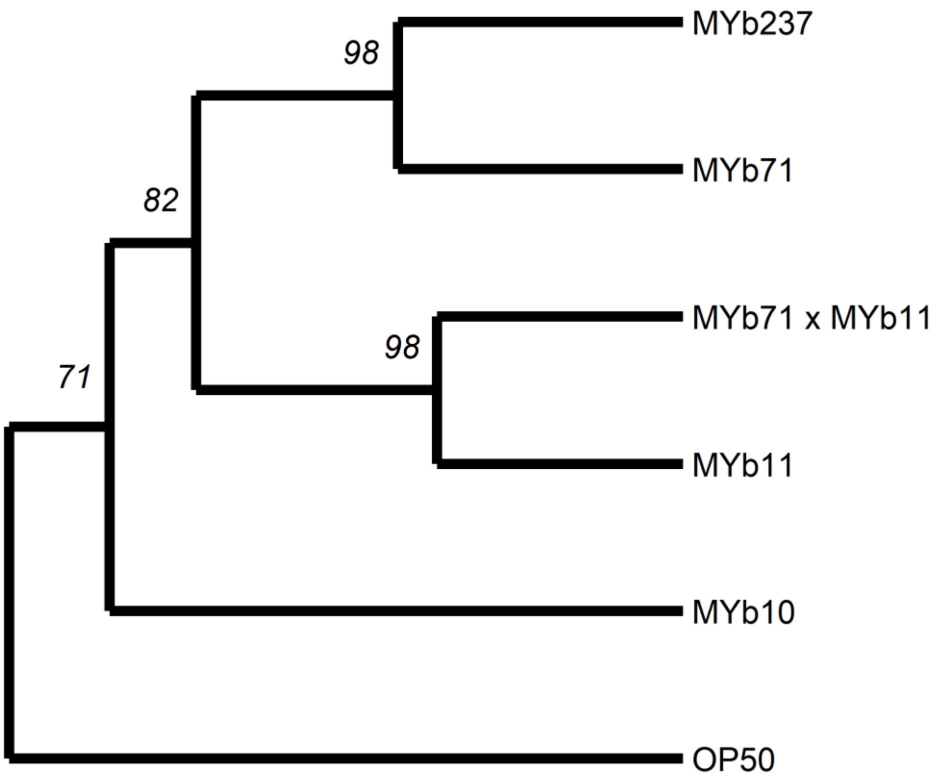


**Supplementary Fig. S4. Hierarchical clustering of strains based on BIOLOG profiles.** Clustering is based on Ward’s algorithm and Euclidean distance measures. Bootstrap support (nboot = 1000) is shown on nodes.

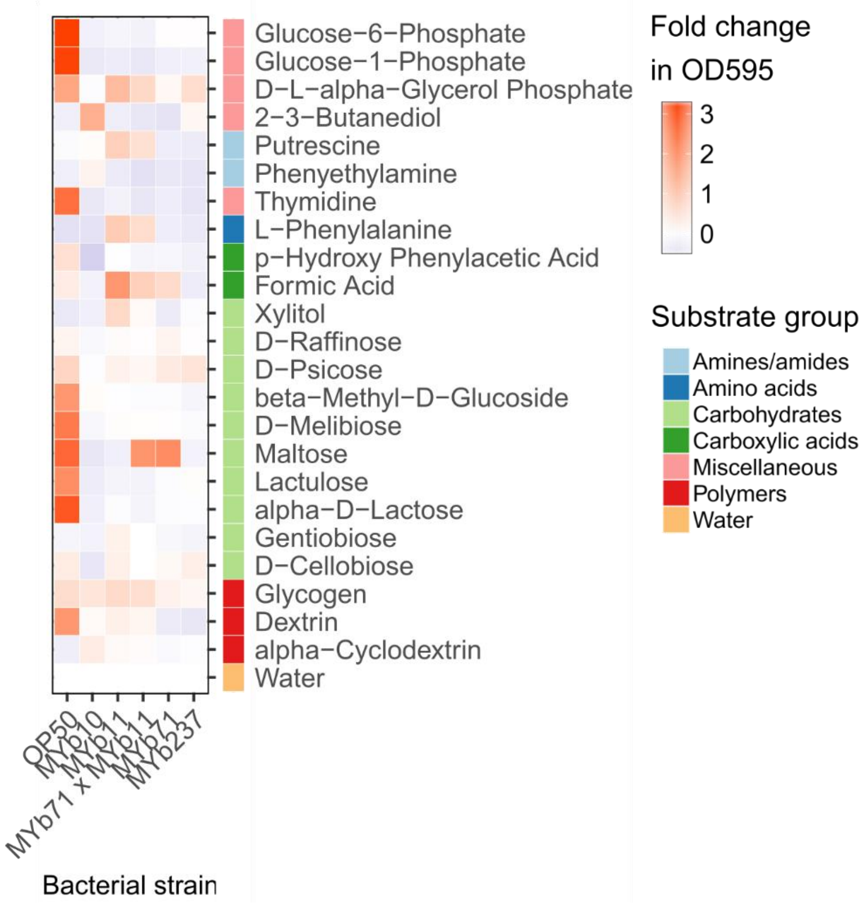


**Supplementary Fig. S5. Cluster 7 of BIOLOG profiling.** Profiles of carbon substrate use of *Acinetobacter* sp. (MYb10), *Pseudomonas lurida* (MYb11), *Ochrobactrum* sp. (MYb71), *Ochrobactrum* sp. (MYb237), and *E. coli* OP50 in BIOLOG GN2 plates over 46 h. The fold-change in indicator dye absorption from 0 to 46 h indicates that the indicated compound is metabolized. K-means clustering (k = 7) of substrates by fold-change highlights metabolic differences between strains. Clusters I - VI are shown in figure 3 in the main text.

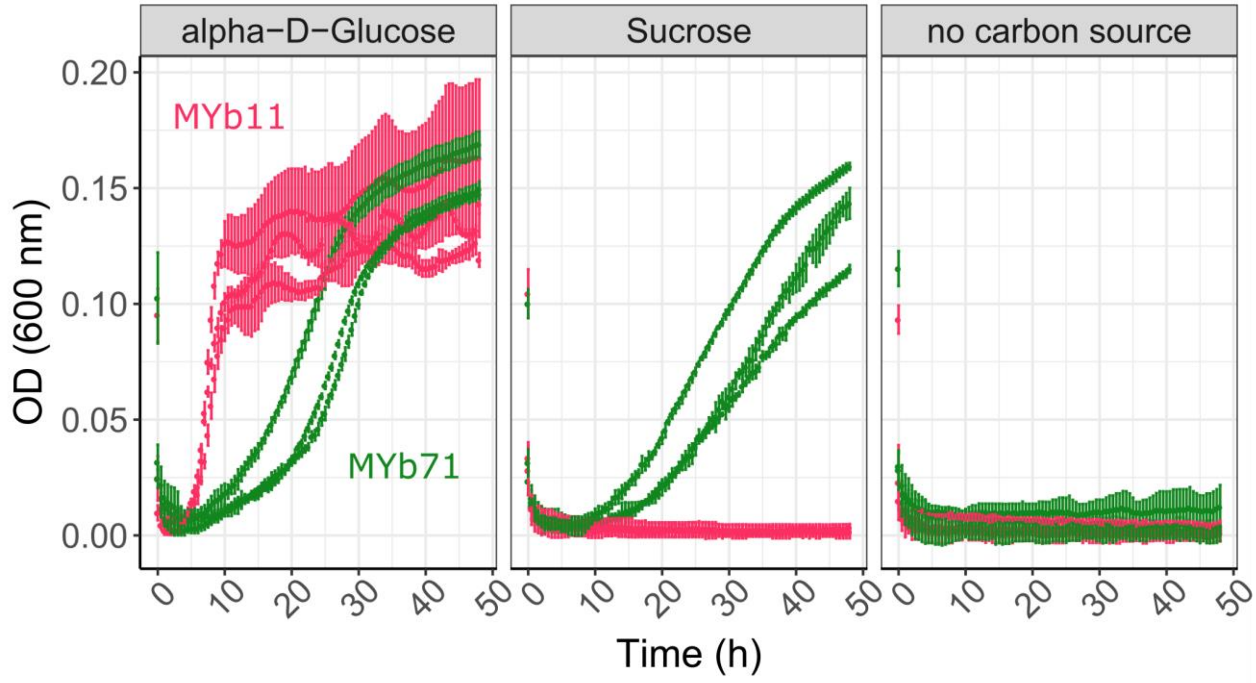


**Supplementary Fig. S6. Culture of MYb11 and MYb71 in defined media with single carbon substrates.** Growth growth of MYb11 and MYb71 in chemically defined media with a single carbon source (i.e., alpha-D-glucose or sucrose), or no-carbon control (over 46 h in 96-well plates).

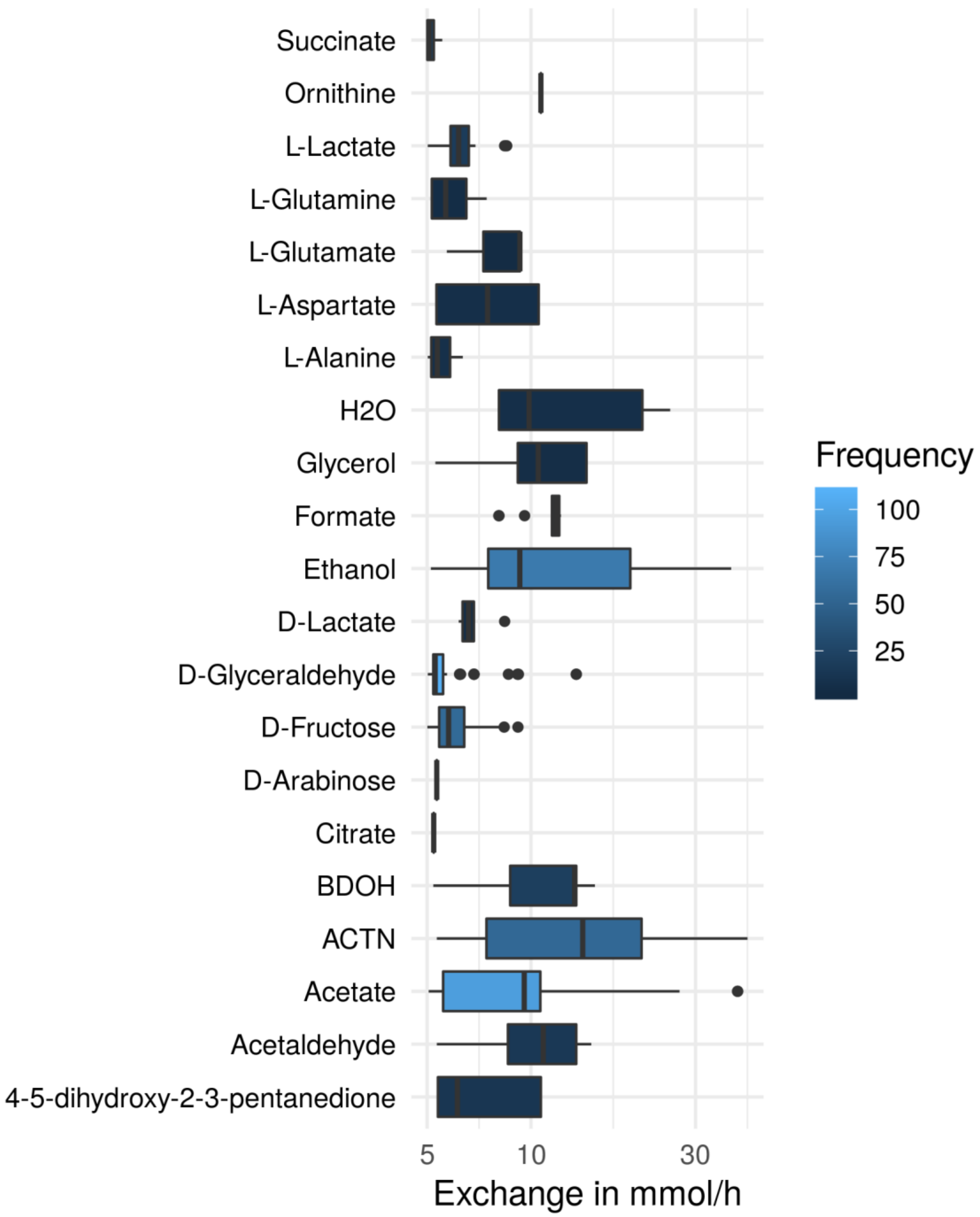


**Supplementary Fig. S7. Metabolites exchanged in *in silico* pairwise interactions on a minimal medium**. Substances which were exchanged between bacterial isolates in simulation of ecological interactions. In a minimal medium, metabolic byproducts could influence the growth rates of organisms. The figure shows metabolites, which were predicted to be exchanged most frequently and in highest quantity.

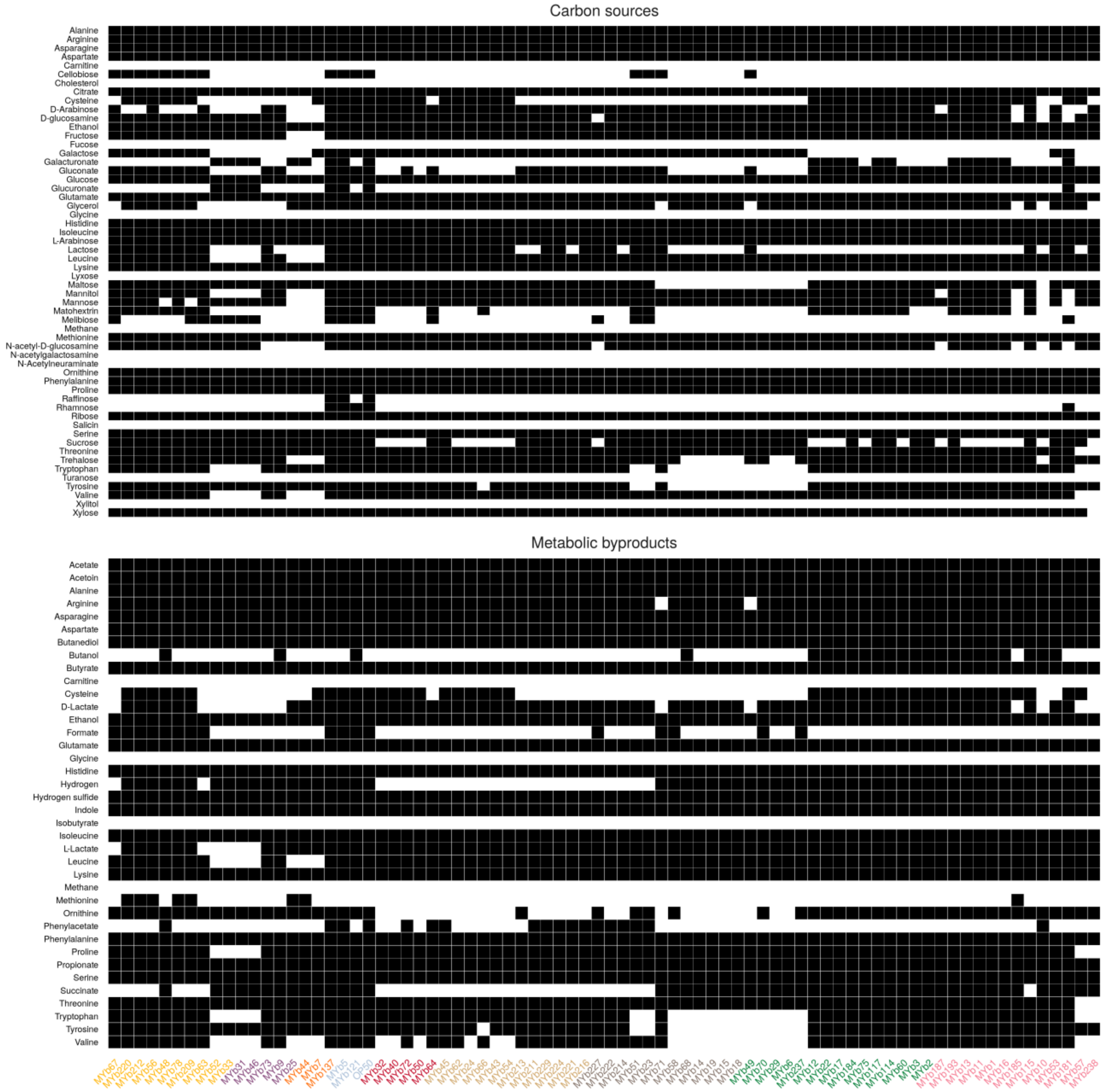


**Supplementary Fig. S9. Prediction of carbon sources and metabolic byproducts**. In simulation of bacteria metabolism, we predicted substances which could be used as carbon sources (left panel) and substances that may be secreted as metabolic byproducts during for example fermentative processes (right panel). Microbiome isolates are given along the x-axis and compounds along the y-axis.

a

**>Sucrose invertase in MYb71 genome**

EIHINGGQHV*----YPNKARRLKFSALLCLCLF*SI*QLSCV
FMILCCGESLIDMLPRETAAG--ETAFQPFAGGSVFNTAIA
LGRLDVPTGFFSGISSDFFGEVLRDNLARSNVDYSFAAIS
DRPTT-LAFVRL-VDGQARYAFYDENTAGRMLTESDMPY-
VDDAIDAMLFGCISLISEPCGSVYEALMT-REAPRRVMFL
DPNIRAGFITDREKHLHRMKRMIALADIVKLSDEDLAWF
GEKGSHDEIAAEWLKLGPKLVVITKGAHGADAYTAKATV
RVPGVKVDVVDTVGAGDTVNAGILASLHNQGLLDKDAL
VELTEDQIHSAVALGVRAAAVTVSRAGANPPW

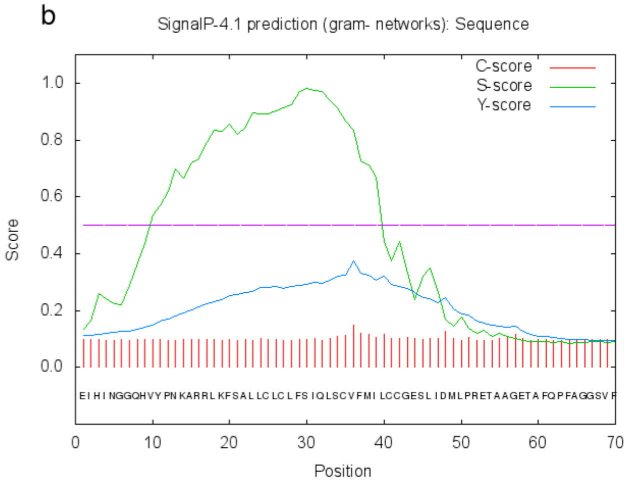


**Supplementary Fig. S10. External sucrose invertase in MYb71.** (a) Amino acid sequence of the sucrose invertase found in the MYb71 genome. (b) We used SignalP to check for secretory signatures. We found that a discrimination score of D = 0.518 (i.e. weighted average of the mean S and the max. Y scores) which indicates a signal peptide (signalP: Name=Sequence SP='YES' Cleavage site between pos. 35 and 36: LSC-VF D=0.518 D-cutoff=0.420 Networks=SignalP-noTM).

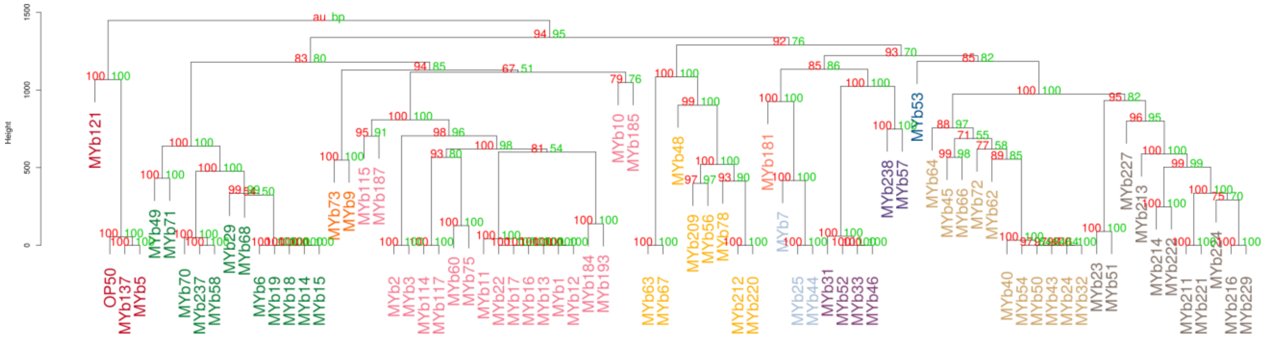


**Supplementary Fig. S11. Hierarchical clustering of metabolic networks based on pathway prediction.** Metabolic networks were clustered according to their pathway completeness score by Euclidean distances and similarity of clusters was estimated by average linkage. The quality of the clustering was tested by multiscale bootstrap resampling. Green values next to branches indicate the bootstrap probability (number of 90 means e.g. that the cluster exists in 90 of 100 runs). In addition to this, the approximately unbiased p-value from multiscale bootstrap is shown in red (see (38) for details).

*Supplementary tables*

**Table S1: Genome characteristics** (Samples_all.xlsx). Genomic overview of all bacterial isolates. Based on sequence identity, the closest related species/strains are shown in column B. Genome and assembly statistics (contigs, N50, GC, ...) can be found in columns G:M. The corresponding bioproject to access data via NCBI is shown in column N.

**Table S2: *In silico* TSB-based medium** (In silico TSB medium.csv). Compounds and maximal uptake rates for Tryptic soy broth (TSB) medium used to simulate bacterial growth.

**Table S3: *In silico* glucose minimal medium with thiamine** (in silico glucose minimal medium with thiamine.csv). Compounds and maximal uptake rates for glucose minimal medium with thiamine used to simulate bacterial growth.

**Table S4: Predicted pathways** (predicted_pathways.xlsx). For each bacterial isolate (column A) the predicted presence (column D and column E for with more conservative bitscore cutoff) for all considered metabolic pathways (column C) is given. Here, a number of 1 means present and 0 means not available. In column G the hierarchy (i.e. subsystem) of the pathway is shown.

**Table S5: Predicted virulence** (Predicted_virulence.xlsx). In this table the presence of virulence traits based on homology with the virulence factor database is shown. In column A the isolates are given and columns B:AV show the presence of virulence factors (1 for true and 0 for false).

**Table S6: Traits with differences in *Ochrobactrum*** (ochrobactrum.xls). Table consists of metabolic pathways and virulence factors which showed to be significantly changed in isolates belonging to the *Ochrobactrum* genus (based on a Wilcoxon signed rank test). For each trait (column B) a FDR corrected P-value (column C) and the mean for Ochrobactrum (column D) and all other isolates (column E) is given.

**Table S7: Experimental phenotypic data: bacterial load and *C. elegans* population growth.** (TableS_phenotypes.xlsx) Sheet 1 shows the number of colony forming units per worm (column D) shown across microbiome isolates (column A) and experimental repetitions of the analysis (i.e., runs; column C). Sheet 2 shows the mean number of worms counted (3 samples counted per population) per population of worms (column C) and standard deviation (column D) on the respective microbiome isolates (column A).

**Table S8: Regression analysis to infer metabolic competences associated with bacterial colonization and host fitness.** (colonization_regression.xlsx) Two regression approaches were used to find traits which were associated with experimental data of bacterial load in *C. elegans* and the fitness of *C. elegans* when the worm was grown together with the bacterial isolates. In this table, the significant/important traits of both approaches are listed.

**Table S9: Traits and scores used to categorize isolates according to adaptive strategies**. (adaptive_strategies.xlsx) Table of metabolic traits and model features associated with stress-tolerating, competitive, or ruderal strategies. For each isolate the scores and classification for each strategy are listed.

*Supplementary data*

**Supplementary data S1. Genome-scale metabolic models of the microbiome of C. elegans.** Zip archive of metabolic models for each isolate in Systems biology markup language (SBML) format. In addition to this, an R file with a list of all models is provided.
